# Contesting the evidence for −1 frameshifting in immune-functioning C-C chemokine receptor 5 (*CCR5*) – the HIV-1 co-receptor

**DOI:** 10.1101/513333

**Authors:** Yousuf A. Khan, Gary Loughran, John F. Atkins

## Abstract

During the decoding of a subset of mRNAs a proportion of ribosomes productively shift to the −1 reading frame at a specific, slippage-prone site^1^. While the great majority of occurrences of the “Programmed −1 Ribosomal Frameshifting” (−1 PRF) involve viruses^2^, other mobile elements or retroelements^1^, a dramatic instance of utilized human cellular −1 frameshifting in a non-retroelement derived mRNA has been reported. The mRNA for which −1 frameshifting was claimed due to a stimulatory pseudoknot is that which encodes immune functioning C-C chemokine receptor 5 (CCR5), the HIV-1 co-receptor^3^. The publication ascribed the importance of the reporter *CCR5* mRNA frameshifting directing translating ribosomes to a premature termination codon leading to mRNA destabilization via the nonsense-mediated decay pathway (NMD), rather than creating a C-terminally unique peptide. The same lab estimates that up to 10% of eukaryotic genes may be regulated by this frameshift-NMD mechanism^4^ and point to the potential for novel therapeutic strategies aimed at treating disease by −1 PRF^5^.

Efficient −1 frameshifting requires a slippery site together with an appropriately positioned stimulatory RNA structure. The reported *CCR5* slip site (U_UUA_AAA where underscores set of the reading frame) has previously been shown to facilitate high levels of −1 frameshifting in vitro when positioned 5’ of a known stimulatory pseudoknot^6^. However, frameshifting efficiencies in vivo could be substantially different and further, at least as of 2016, there were no known virus frameshift sites with the reported *CCR5* shift site^1^.

We challenge the reported results by showing that frameshifting at the relevant *CCR5* sequence cassette does not occur above background levels and that the claimed pertinent sequence is not as distinctively conserved as previously presented. Lastly, we demonstrate that previously published retroelement derived human frameshift signals do indeed harbor frameshift activity.

Using BLAST, we first identified orthologues of human *CCR5* in the NCBI RefSeq database. Sequences were aligned and, on examining the frameshift region, we observed substitutions in the slippery site sequence which would likely decrease or eliminate frameshifting in related organisms. Indeed, none of the nucleotides in the slippery site are conserved across vertebrates (Figure 1). The predicted frameshift site in *CCR5* was previously reported as most highly conserved in primates, except for *Lemur catta*^3^. Primate *CCR5* CDS sequences were aligned and nucleotide conservation was measured at single nucleotide resolution (Figure 2A) and smoothed in a 10 (Figure 2B) nucleotide sliding window. This analysis showed that the entire CDS, not just the predicted frameshift site, is highly conserved among primates. Thus, the observation that the frameshift site is mostly conserved in primates does not indicate that the (putative) ability to frameshift is subject to purifying selection.

**Figure 1:**
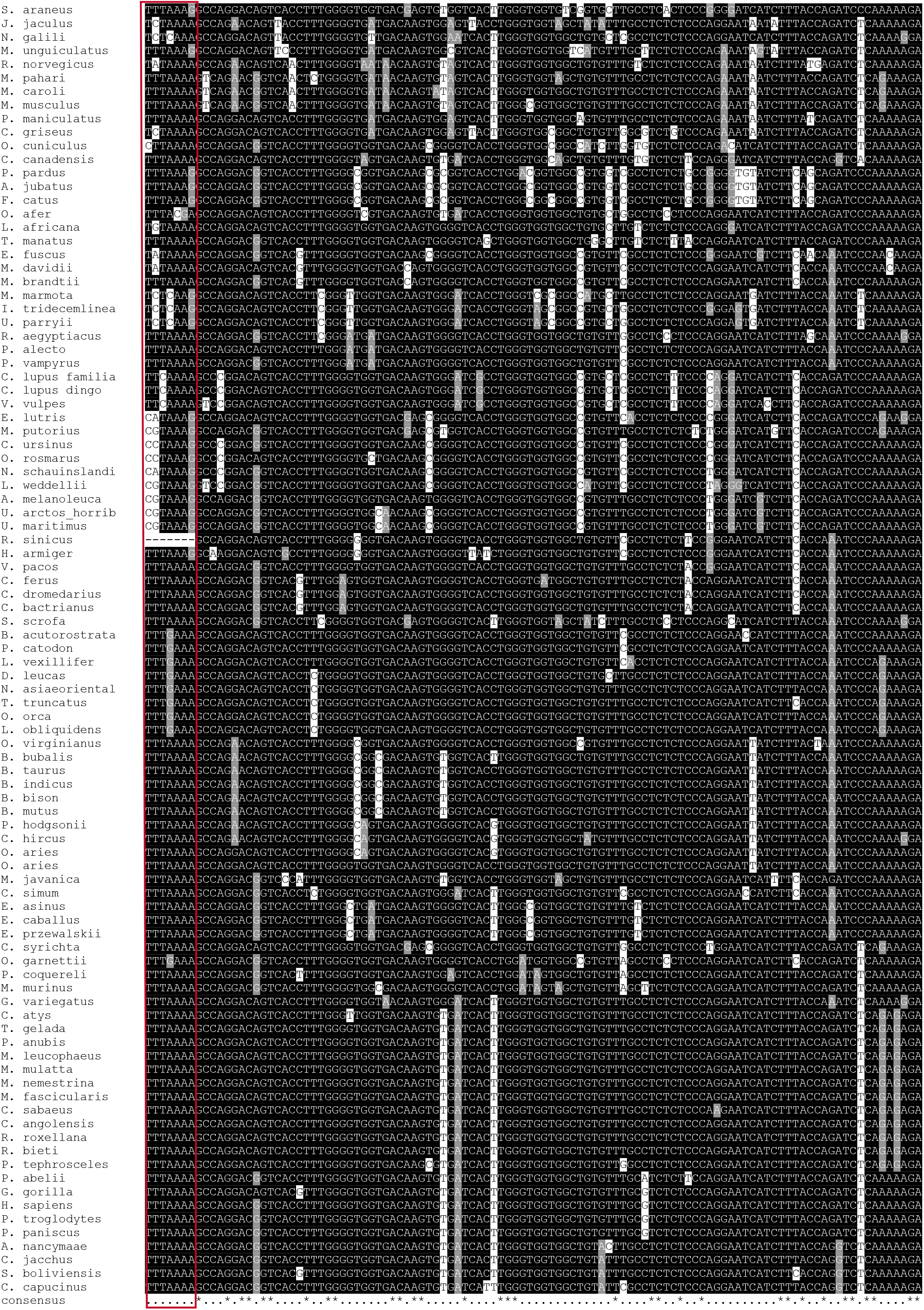
Alignment of putative frameshift site: Sequence alignment of the *CCR5* CDS regions from BLAST hits to human *CCR5*, trimmed to only include the frameshift region. The red box indicates the putative slippery site.

**Figure 2:**
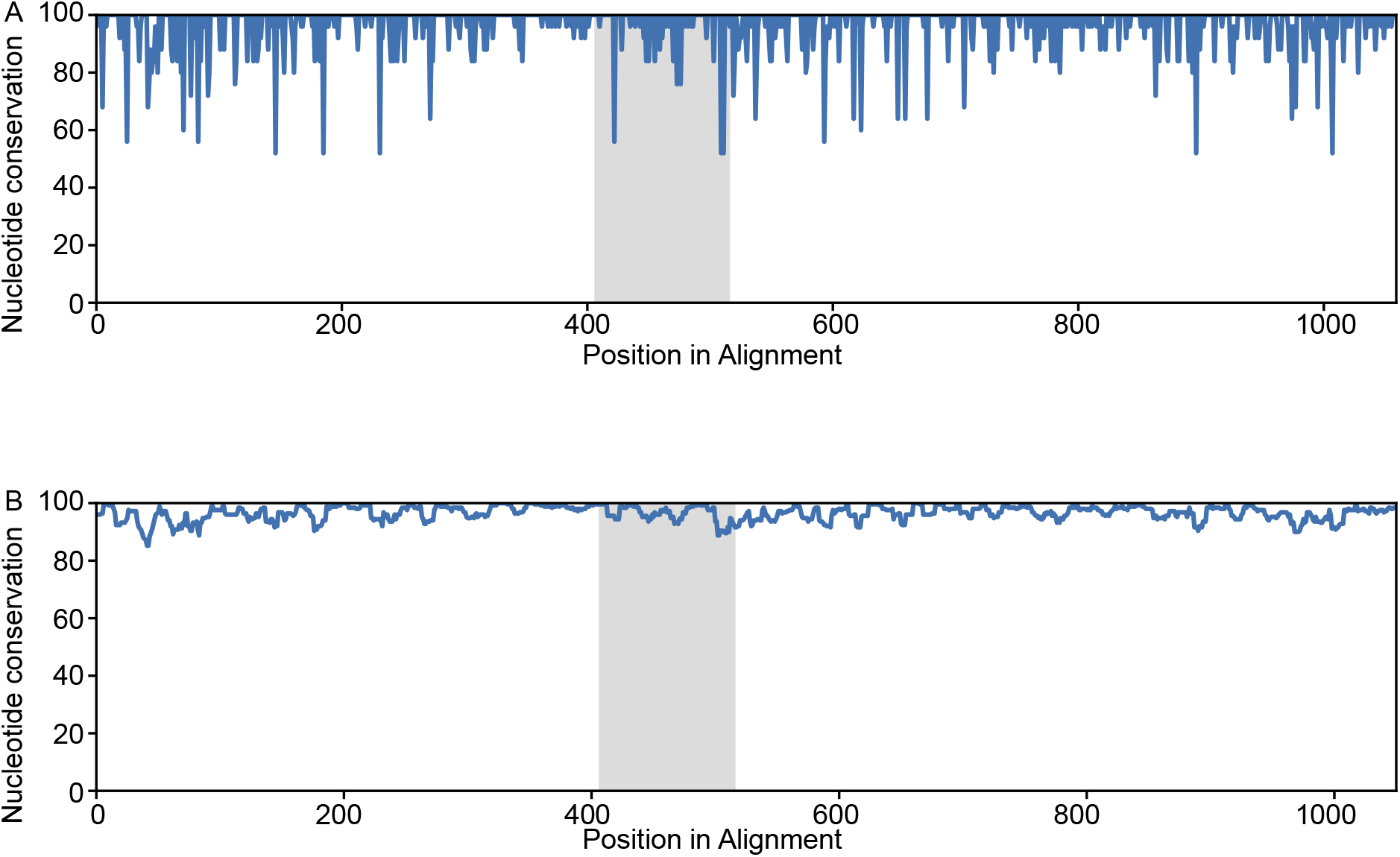
Sliding window conservation analysis of the *CCR5* CDS: **a.** Single nucleotide resolution in the *CCR5* CDS for primates only, excluding *Lemur catta*. **b.** 10-nucleotide sliding window of conservation in the *CCR5* CDS for primates only, excluding *Lemur catta*. The grey highlight represents the *CCR5* predicted frameshift region.

Dual-luciferase vectors in which the test cassette is sandwiched between and fused to both Firefly and Renilla luciferase encoding sequences, have been widely used for assaying translational recoding events (Figure 3A). The frameshifting efficiency is calculated by dividing Firefly luciferase activity by Renilla luciferase activity. This ratio is then normalized to an in-frame control vector in which Renilla and Firefly luciferases are both situated in the zero-frame to account for differences in activity between the two enzymes (a ratio of a ratio) (Figure 3B).

**Figure 3:**
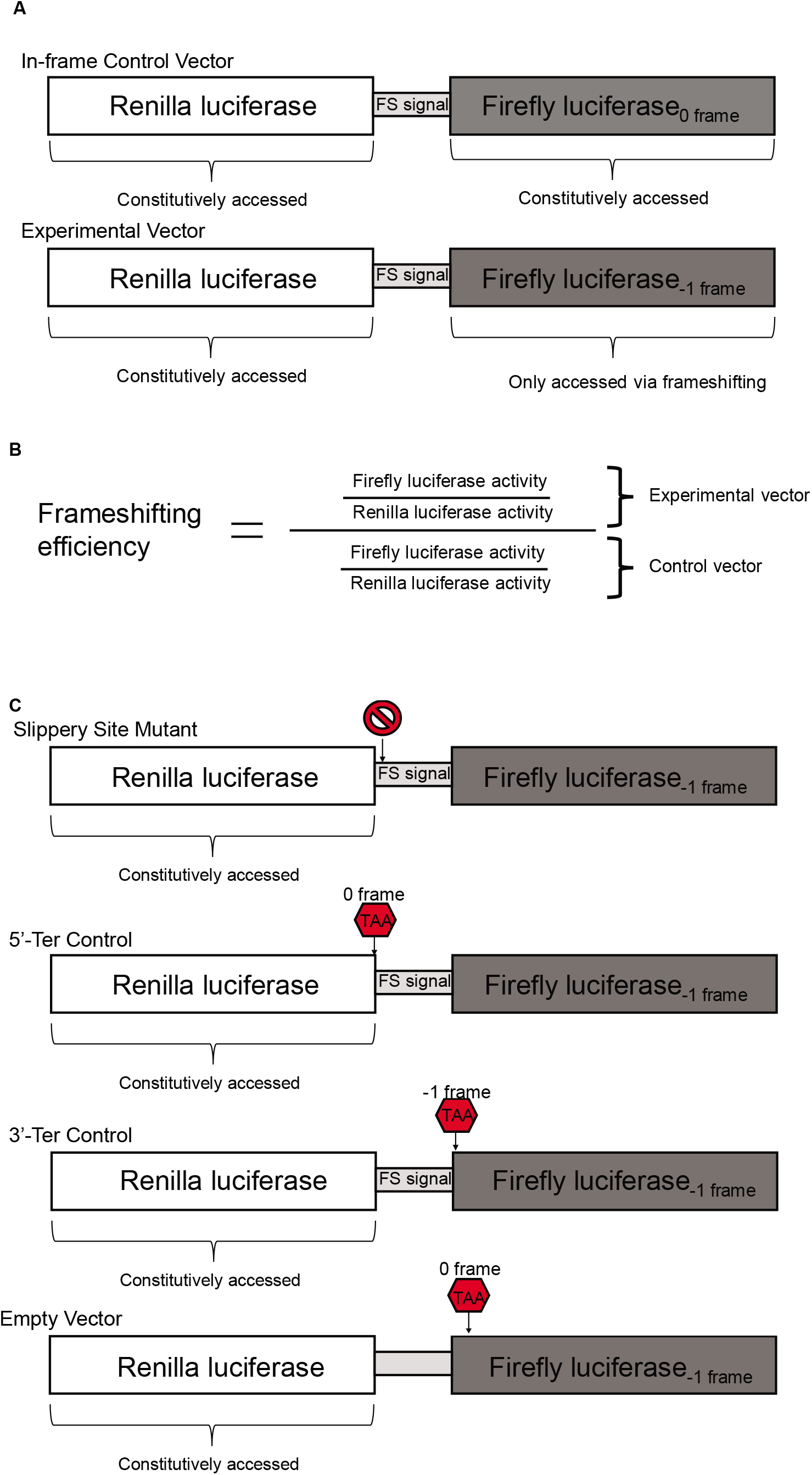
Schematic of the dual luciferase vector: **a.** Graphical representation of experimental and in-frame control for dual luciferase assays. **b.** Mathematical formula for calculating rates of ribosomal frameshifting. **c.** Graphical representation of negative controls.

In nearly all cases the fused test encoded polypeptide has only a small or no effect on the upstream encoded luciferase activity. However, in one case luciferase activity was anomalously low (unpublished) and we developed a variant vector system (pSGDluc^7^) containing a StopGo sequence directly 3’ of the first and immediately 5’ of the second luciferase encoding sequence. Subsequently we made another variant by removing a potential splice site, pSGDlucV3.0 (deposited with Addgene). Thus in earlier constructs, where the insert-encoded peptides were appended to the C-terminus of Renilla luciferase, the Renilla luciferase activity could drop – particularly for the short zero-frame encoded peptide, causing an artificial inflation of translation recoding rates. This problem has been overcome by inserting flanking StopGo sequences that result in co-translational excision of the insert-encoded peptides, thus releasing independent Renilla and Firefly proteins that are identical in sequence for both the test and in-frame control constructs. The publication on *CCR5* frameshifting utilized the very first iteration of a dual luciferase vector that we developed in 1998^8^.

We recloned the same *CCR5* frameshift sequence tested in the original article into pSGDlucV3.0. For *CCR5* we used negative controls comparable to those in the original article, which include a slip site mutant (UUUAAAA → GCGCGCG for **Y.A.K.**, UUUAAAA → AUUCAAA for **G.L.**), a 5`-termination control (also known as a premature termination codon control or PTC control) and a 3`-termination control (also known as an out-of-frame stop codon control or OOF control). Lastly, we included measurements from the empty vector alone (Figure 3C). All four of these controls are meant to establish a baseline for background levels of luminescence

The dual luciferase assays were carried out in the same cell line used in the original article, HeLa. Renilla luciferase, the reference luciferase, is stable for all the constructs (Figure 4A). Similarly Firefly luciferase activities for the in-frame controls of *CCR5 and* a HIV-1 control are consistent as well, indicating that when forced to express Firefly luciferase, both constructs produce proteins with similar activity. The Firefly luciferase activity for the WT HIV-1 frameshift construct is >40 fold higher than that of WT *CCR5* or any of its negative controls. Furthermore, the Firefly luciferase activity for *CCR5* and all its negative controls are all comparable to the activity of the empty vector control, which is the baseline for background levels of Firefly luciferase activity (Figure 4B). After normalizing the luciferase activities for the test vectors to their respective in-frame control vectors, we find that HIV-1 shows frameshift rates that are comparable to previously reported findings in HeLa cells^9^. However, *CCR5* WT frameshifting is ~0.15%, indistinguishable from its negative controls, indicating that *CCR5* does not frameshift above background at our limit of sensitivity and is certainly far less than the previously reported level of 9-11%^3^. In fact, *CCR5* SSM demonstrates slightly higher rates of FS than the WT, indicating that translational recoding values this low are indicative of background (Figure 4C). These experiments were repeated in HEK293T cells to control for cell-line specific results and we found that the results followed identical trends to those obtained in HeLa cells (Figure 4D-F).

**Figure 4:**
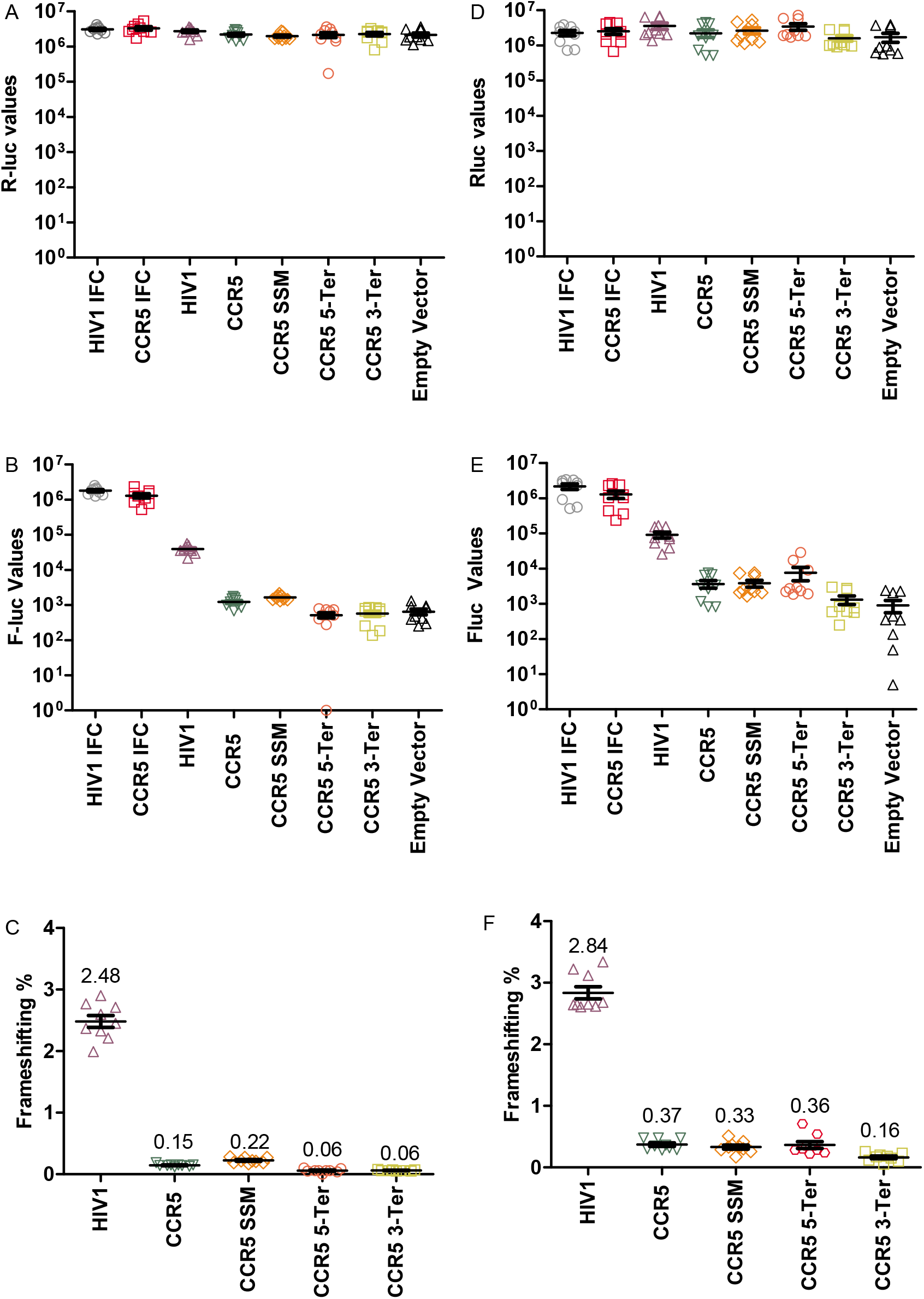
The *CCR5* sequence does not promote efficient frameshifting above its negative controls: **a.** Absolute Renilla luciferase values for all experiments performed in HeLa cells. **b.** Absolute Firefly luciferase values for all experiments performed in HeLa cells. **c.** Frameshifting efficiency for all experiments performed in HeLa cells, normalized to either the HIV-1 or *CCR5* in-frame control vector as appropriate. **d.** Absolute Renilla luciferase values for all experiments performed in HEK293T cells. **e.** Absolute Firefly luciferase values for all experiments performed in HEK293T cells. **f.** Frameshifting efficiency for all experiments performed in HEK293T cells, normalized to either the HIV-1 or *CCR5* in-frame control vector as appropriate. Every data point represents an independently transfected well of cells and error bars represent standard deviations.

To ensure that these results were repeatable, the cloning and assaying of *CCR5* frameshifting was completed independently by a different laboratory (**G.L., J.F.A.**), which used modestly different cloning and assaying strategies. When utilizing both HeLa and HEK293T cells, we found that *CCR5* frameshifted at an insignificant rate as well (Figure 5A-F).

**Figure 5:**
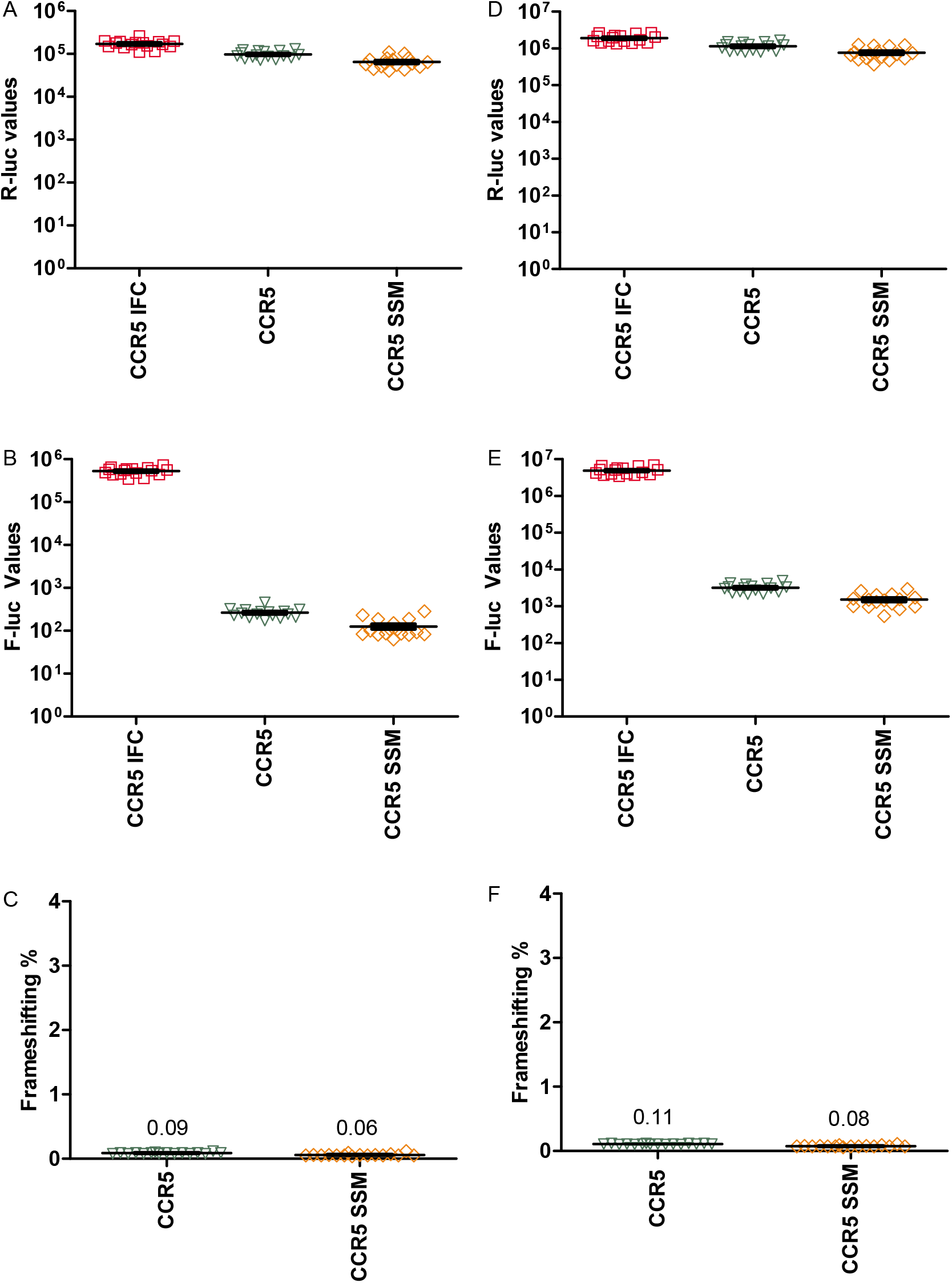
Independent confirmation that the *CCR5* sequence does not promote frameshifting above its negative controls. **a.** Absolute Renilla luciferase values for all experiments performed in HeLa cells. **b.** Absolute Firefly luciferase values for all experiments performed in HeLa cells. **c.** Frameshifting efficiency for all experiments performed in HeLa cells, normalized to the *CCR5* in-frame control vector. **d.** Absolute Renilla luciferase values for all experiments performed in HEK293T cells. **e.** Absolute Firefly luciferase values for all experiments performed in HEK293T cells. **f.** Frameshifting efficiency for all experiments performed in HEK293T cells, normalized to the *CCR5* in-frame control vector. Every data point represents an independently transfected well of cells and error bars represent standard deviations.

Retrotransposons and retroviruses share a common origin. Ribosomal frameshifting has been well documented in viruses and thus it stands to reason that retrotransposons and/or derivatives of retrotransposons in the human genome could frameshift. Three derivatives of retrotransposons in the human genome, *PNMA3 (Ma3), PNMA5 (Ma5)*, and *PEG10*, have been previously demonstrated to frameshift in-vivo and in-vitro^1^. All three encode zero-frame proteins with homology to the viral protein *gag* and their trans-frame protein have similarity to *gag-pol*. All three share a slippery site common with some viruses in the *Retroviridae* family, GGGAAAC. These sequences were assayed in both HeLa and HEK293T cells (Figure 6). Although the rates of −1 PRF were slightly different from previously reported, all three signals stimulated frameshifting far above background. These results further confirm the fidelity of the pSGDlucV3.0 vector for assaying −1 PRF, already seen for other translational recoding events^7^.

**Figure 6:**
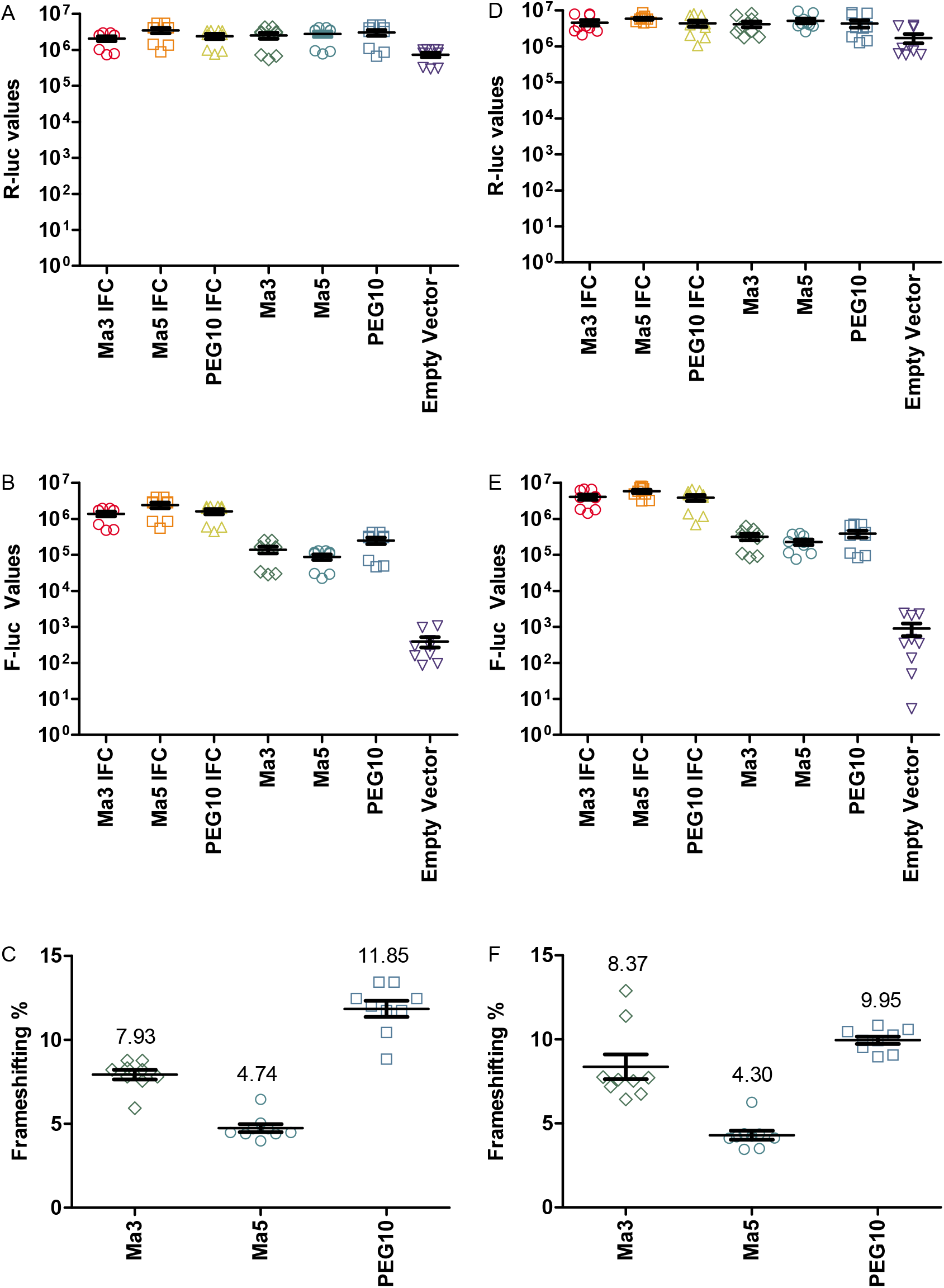
Previously validated retrotransposon-derived frameshift signals in human still demonstrate efficient frameshifting. **a.** Absolute Renilla luciferase values for all experiments performed in HeLa cells. **b.** Absolute Firefly luciferase values for all experiments performed in HeLa cells. **c.** Frameshifting efficiency for all experiments performed in HeLa cells, normalized to the respective in-frame controls *Ma3, Ma5*, or *PEG10*, as appropriate. **d.** Absolute Renilla luciferase values for all experiments performed in HEK293T cells. **e.** Absolute Firefly luciferase values for all experiments performed in HEK293T cells. **f.** Frameshifting efficiency for all experiments performed in HEK293T cells, normalized to the respective in-frame controls *Ma3, Ma5*, or *PEG10*, as appropriate.

In conclusion, using an improved reporter assay system, we find no evidence of −1 frameshifting above background levels on the CCR5 mRNA. Additionally, a lack of statistically significant purifying selection acting on the slip site (a hallmark for functionally relevant frameshift signals) supports the argument that *CCR5* does not contain a biologically important frameshift signal. There are several possible explanations for the discrepancy between the reported *CCR5* results and the results presented here but the most obvious one is the standard decoding of a significant percentage of mRNAs generated by cryptic splicing.

## Methods

### Alignment of *CCR5* slippery sites

The NCBI mRNA and RefSeq BLAST databases were searched with blastn using the human CCR5 mRNA (accession NM_000579.3) as query on December 4th, 2018 using default parameters, with the number of hits expanded from 100 to 500. The best sequence per organism, as determined by ‘Max Score’, was taken up until 70% query cover. These sequences were aligned with MUSCLE. This alignment was trimmed to only include the putative frameshift region of CCR5 that was generated by code written in Python 3. The alignment image was generated using the box-shade server (https://embnet.vital-it.ch/software/BOX_form.html).

### *CCR5* Primate CDS analysis

Of the sequences obtained in the ortholog search, only those belonging to the taxonomic descriptor of ‘primate’ (taxid 9443) were used. *Lemur catta* sequences were ignored, since the original manuscript states that the *CCR5* frameshift signal is not conserved in *Lemur catta*. All CDS sequences were aligned with MUSCLE and then either at single nucleotide resolution or a smoothed 10 nucleotide window plotted in Python 3.

### Vector Construction

pSGDlucV3.0 was generated by introducing silent mutations into the Renilla coding sequence to disrupt two potential donor splice sites (TGGgtaagt).

For Figures 4 and 6 (**Y.A.K.**) the vector was digested with *Psp*XI and *Bgl*-II. Overlapping oligonucleotides for insert sequences were ordered from Sigma Aldrich (**Supplementary Table 1**), resuspended at 100 μM, diluted to 10 μM, and then mixed together at a 1:100 dilution. This 1:100 dilution of the forward and reverse oligonucleotide was heated to 95°C. 2 μL of this mix was ligated into 40 ng of linearized backbone vector using NEB T4 ligase.

For Figure 5 (**G.L.**) The vector was digested with *Psp*XI and *Bgl*-II. Using forward and reverse PCR primers (**Supplementary Table 1**), the *CCR5* WT, SSM and IFC were all amplified from genomic DNA with enzyme overhangs. These PCR products were subsequently digested and ligated into pSGDlucV3.0.

### Transfections and Dual Luciferase Assays

For Figures 4 and 6 (**Y.A.K.**) HeLa and HEK293T cells were reverse transfected with 100 ng of plasmid using 0.2 μL Lipofectamine 2000 in a 96-well format. Transfection mix was added to 30,000 cells and incubated for 1-2 minutes at room temperature, before adding Fetal Calf Serum. The next day, media was removed from cells and 100 μL of fresh 1X passive lysis buffer (Promega) was added to each well. Samples were frozen and thawed before luciferase activity was measured by adding 20 μL of lysate and 20 μL of each reagent, as per the Promega protocol.

For Figure 5 (**G.L.**) HeLa and HEK293T cells were reverse transfected with 25 ng of plasmid using 0.2 μL Lipofectamine 2000 in a half-area 96-well format. Transfection mix was added to 40,000 cells. The next day, transfected cells were lysed in 12.6 μL of 1X passive lysis buffer (Promega) and luciferase activity was measured by injection of 20 μL of each reagent.

Background level readings (measured from water transfected cells) were subtracted from all Firefly and Renilla luciferase readings. Frameshift efficiencies were calculated by dividing Firefly luciferase values by Renilla luciferase values and then dividing the relative ratios by the average F-luc/R-luc ratio of the in-frame control.

## Author Contributions

Y.A.K. conceived and coordinated the project. G.L. established the pSGDlucV3.0 vector used in this study. Y.A.K. performed the experiments and analyzed the data shown in Figures 4 and 6 whereas G.L. independently conducted experiments and analyzed the data shown in Figure 5. Y.A.K. performed the comparative phylogenetic analyses shown in Figures 1 and 2. Y.A.K. wrote the manuscript and created the figures. All authors edited and approved the manuscript.

## Acknowledgements

This work was supported by grants from Science Foundation Ireland (12/IP/1492 and 13/1A/1853) to J.F.A. Y.A.K. was supported by an award from the Winston Churchill Foundation of the United States of America.

---

**Supplementary Table 1.**
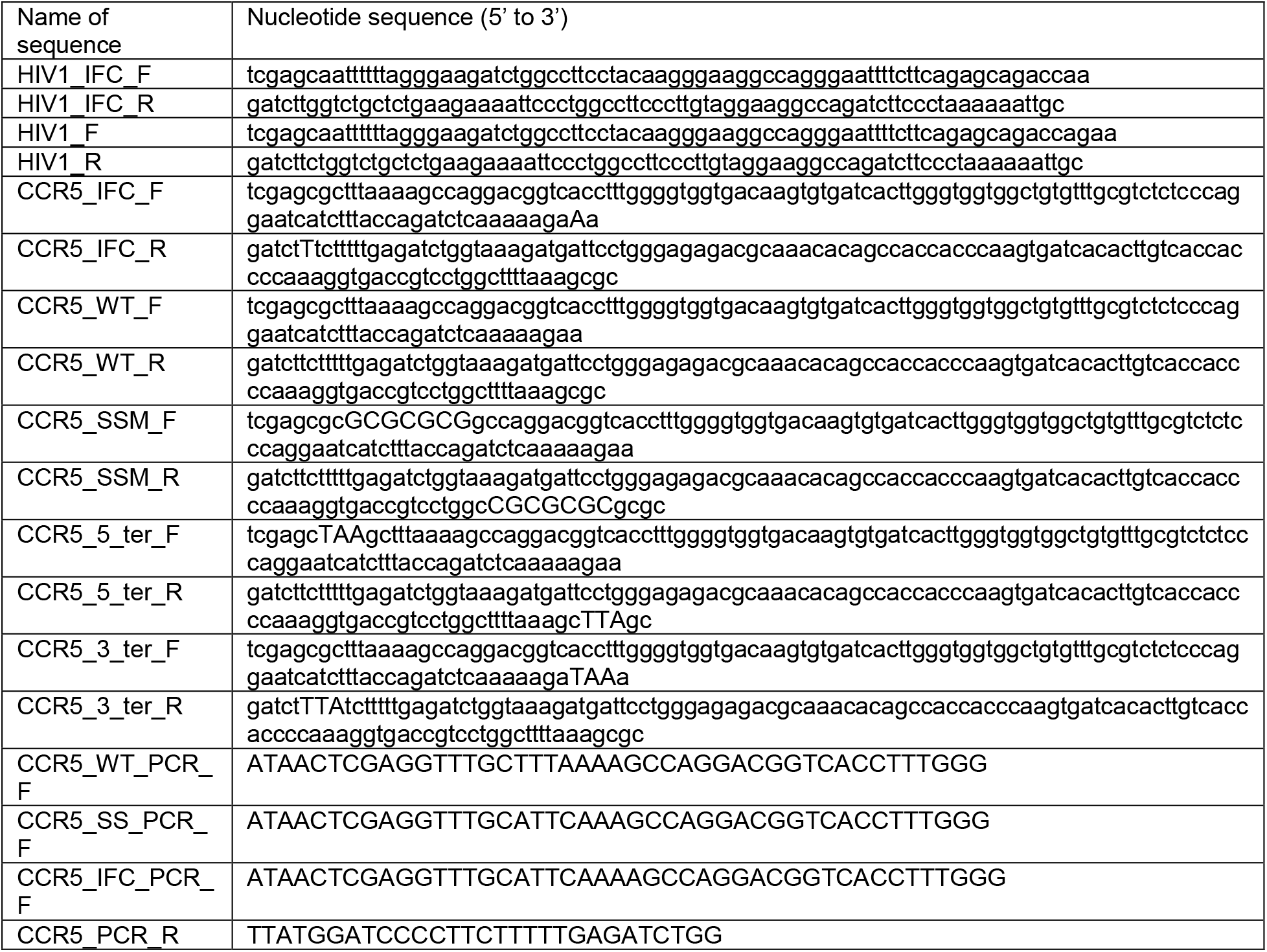

**Supplementary Table 2 -.** Accessions

|  |
| --- |
| NM_001100168 |
| NM_001009046 |
| NM_001135497 |
| XM_003818462 |
| XM_004034017 |
| NM_001042773 |
| NM_001309022 |
| XM_005546902 |
| XM_025376087 |
| XM_023231324 |
| XM_011939245 |
| XM_010374788 |
| XM_017855753 |
| XM_009200976 |
| XM_011981068 |
| XM_007983952 |
| XM_012033360 |
| XM_012470795 |
| NM_001257216 |
| XM_012638782 |
| XM_003926266 |
| XM_008071218 |
| XM_017521787 |
| XM_012770304 |
| XM_020916150 |
| NM_001011672 |
| XM_013981569 |
| XM_006173924 |
| XM_010997075 |
| XM_010967349 |
| XM_008582128 |
| XM_025884850 |
| XM_017666109 |
| XM_006071554 |
| XM_003794523 |
| XM_019985017 |
| XM_010862398 |
| XM_005900894 |
| XM_005955139 |
| XM_007168860 |
| XM_006200721 |
| XM_024123986 |
| XM_004399154 |
| XM_021683054 |
| XM_015489371 |

